# The role of competition versus cooperation in microbial community coalescence

**DOI:** 10.1101/2021.04.18.440290

**Authors:** Pablo Lechón, Tom Clegg, Jacob Cook, Thomas P. Smith, Samraat Pawar

**Affiliations:** Department of Ecology & Evolution, University of Chicago, Chicago, IL, USA; Department of Life Sciences, Imperial College London, Silwood Park, Ascot, UK; Centre for Integrative Systems Biology and Bioinformatics, Imperial College London, UK

## Abstract

New microbial communities often arise through the mixing of two or more separately assembled parent communities, a phenomenon that has been termed “community coalescence”. Understanding how the interaction structures of complex parent communities determine the outcomes of coalescence events is an important challenge. While recent work has begun to elucidate the role of competition in coalescence, that of cooperation, a key interaction type commonly seen in microbial communities, is still largely unknown. Here, using a general consumer-resource model, we study the combined effects of competitive and cooperative interactions on the outcomes of coalescence events. In order to do so, we simulate coalescence events between pairs of communities with different degrees of competition for shared carbon resources and cooperation through cross-feeding on leaked metabolic by-products (facilitation). We also study how structural and functional properties of post-coalescence communities evolve when they are subjected to repeated coalescence events. We find that in coalescence events, the less competitive and more cooperative parent communities contribute a higher proportion of species to the new community, because this endows superior ability to deplete resources and resist invasions. Consequently, when a community is subjected to repeated coalescence events, it gradually evolves towards being less competitive and more cooperative, as well as more species rich, robust and efficient in resource use. Encounters between microbial communities are becoming increasingly frequent as a result of anthropogenic environmental change, and there is great interest in how the coalescence of microbial communities affects environmental and human health. Our study provides new insights into the mechanisms behind microbial community coalescence, and a framework to predict outcomes based on the interaction structures of parent communities.

**Author summary:** In nature, new microbial communities often arise from the fusion of whole, previously separate communities (community coalescence). Despite the crucial role that interactions among microbes play in the dynamics of complex communities, our ability to predict how these affect the outcomes of coalescence events remains limited. Here, using a general mathematical model, we study how the structure of species interactions confers an advantage upon a microbial community when it encounters another, and how communities evolve after undergoing repeated coalescence events. We find that competitive interactions between species preclude their survival upon a coalescence event, while cooperative interactions are advantageous for post-coalescence survival. Furthermore, after a community is exposed to many coalescence events, the remaining species become less competitive and more cooperative. Ultimately, this drives the community evolution, yielding post-coalescence communities that are more species-rich, productive, and resistant to invasions. There are many potential environmental and health implications of microbial community coalescence, which will benefit from the theoretical insights that we offer here about the fundamental mechanisms underlying this phenomenon.

## Introduction

Microbial communities are widespread throughout our planet [1], from the the human gut to the deep ocean, and play a critical role in natural processes ranging from animal development and host health [2, 3] to biogeochemical cycles [4]. These communities are very complex, typically harbouring hundreds of species [5], making them hard to characterize. Recently, DNA sequencing has allowed high-resolution mapping of these communities, opening a niche for theoreticians and experimentalists to collaboratively decipher their complexity and assembly [6–11, 13].

Entire microbial communities are often displaced over space and come into contact with each other due to physical (e.g., dispersal by wind or water) and biological (e.g., animal-animal or animal-plant interactions, and leaves falling to the ground) factors [14–17]. The process by which two or more communities that were previously separated join and reassemble into a new community has been termed community coalescence [18]. Although microbial community coalescence is likely to be common, the effects of both intrinsic and extrinsic factors on the outcomes of such events remains poorly understood [19].Among extrinsic factors, resource availability, immigration rate of new species, and environmental conditions (especially, pH, temperature, and humidity) are likely to be crucial [20–22]. Among intrinsic factors, the role of functional and taxonomic composition and the inter-species interaction structures of parent communities are expected to be particularly important [20, 21]. We focus on the role of species interactions on community coalescence in this study.

Early mathematical models suggested that in encounters between animal and plants communities, species in one community are more likely to drive those in the other extinct (community dominance) [23, 24]. This was explained as being the result of the fact that communities are a non-random collection of species assembled through a shared history of competitive exclusion, and therefore act as coordinated entities. Recent theoretical work [25] has more rigorously established this for microbial community coalescence events, showing that the dominant community will be the one more capable of depleting all resources simultaneously. Overall, these findings suggest that communities arising from competitive species sorting exhibit sufficient “cohesion” to prevent invasions by members of other communities [26, 27].

However, empirical support for the role of competition alone in coalescence outcomes is circumstantial, and the role of cooperation, which is commonly observed in microbial communities, is yet to be addressed theoretically. For example, during coalescence in methanogenic communities, “cohesive” units of taxa from the community with the most efficient resource use are co-selected [22]; and in aerobic bacterial communities, the invasion success of a given taxon is determined by its community members as a result of collective consumer-resource interactions and metabolic feedbacks between microbial growth and the environment [28]. Nonetheless, neither of these studies addressed the role of competition and cooperation in shaping coalescence success. Yet, these microbial communities exhibit cooperation through a typically dense cross-feeding network, where leaked metabolic by-products of one species are shared as public goods across the entire community [29–31]. Indeed, several studies have suggested that a combination of competitive and cooperative interactions may determine the outcome of coalescence in microbial communities [21, 32, 33].

Here, we focus on the relative importance of competition and cooperation in community coalescence. We use a general consumer resource model that includes cross-feeding to assemble complex microbial communities having different degrees of competition and cooperation. We focus on determining the relative importance of the two types of interactions on outcomes of coalescence events, as well as the subsequent evolution of the structural and functional properties of coalesced communities.

## Methods

### Mathematical model

Our mathematical model for the microbial community dynamics is based on the work of Marsland et al. [6] (see Supporting text section 1, and Fig 1):

**Fig 1.**
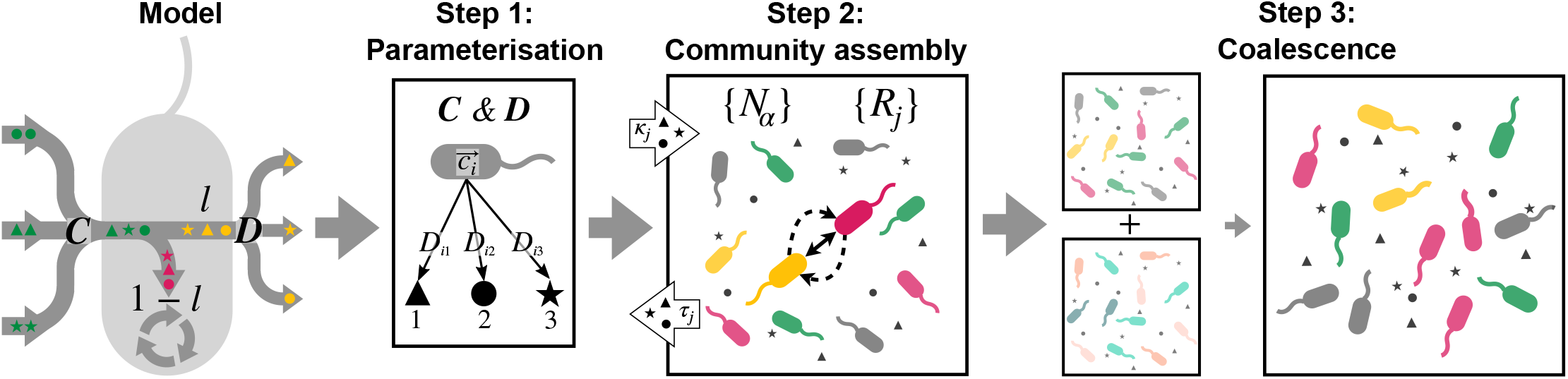
Overview of the coalescence modelling methodology. **Step 1**. The matrix of resource preferences (*C*) and the metabolic matrix (*D*) are sampled for each community. Black polygons are different resource types. **Step 2**. Dynamics of the system are allowed to play out (Eqs 1) until a locally stable equilibrium point is reached. Species composition and abundance, along with community-level competition 𝒞 (solid bidirectional arrows, Eq 3), and facilitation ℱ (dashed unidirectional arrows, Eq 4) are measured in assembled communities. **Step 3**. A pair of the assembled parent communities are mixed, and the resulting community integrated to steady state. For the random and recursive coalescence procedures, the contribution of each parent community to the final mix is analyzed (*S*_1,2_, Eq 5) as a function of their interaction structures (𝒞_1,2_ and ℱ_1,2_) before they coalesced. In the case of the serial coalescence procedure, the properties of the resident community ℛ are tracked after each coalescence exposure.

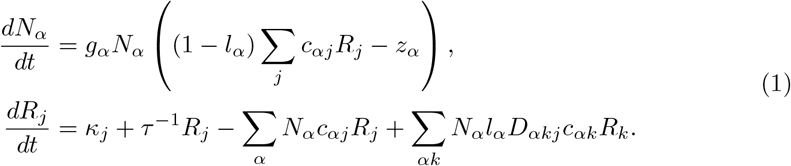

Here, *N*_*α*_ (*α* = 1, …, *s*) and *R*_*j*_ (*j* = 1, …, *m*) are the biomass abundance of the *α*^*th*^ microbial (e.g., bacterial) species and the concentration of the *j*^*th*^ resource (e.g., carbon substrate). The growth of species *α* is determined by the resources it harvests minus the cost of maintenance (two terms in the brackets). Resource uptake depends on the resource concentration in the environment *R*_*j*_, and on the uptake rate of species *α*, here assumed to be binary (*j* (*c*_*αj*_ = 1 or *c*_*αj*_ = 0). The leakage term *l*_*α*_ determines the proportion of this uptake that species *α* releases back into the environment as metabolic by-products, with the remainder (1 − *l*_*α*_) being allocated to growth. The uptake that remains after subtracting a maintenance cost (*z*_*α*_) is transformed into biomass with a proportionality constant of *g*_*α*_, the value of which does not affect the results presented here.

The change in the concentration of resources in the environment (second line in Eq 1) is determined by four terms. The first and second terms represent the external supply and dilution of resource *j*, which give the rates at which the *j*^*th*^ resource enters and leaves the system. The third term is the uptake of the *j*^*th*^ resource from the environment, summed across all *s* consumers, and the fourth term represents resources entering the environmental pool via leakage of metabolic by-products. By-product leakage of species *α* is determined by the metabolic matrix *D*_*α*_ (or the “stoichiometric” matrix; [6]), with *D*_*αjk*_ representing the leaked proportion of resource *j* that is transformed into resource *k* by species *α*. Energy conservation dictates that *D*_*α*_ is a row stochastic matrix, meaning that its rows sum to 1. Note that in this model, rates of metabolic by-product formation are dependent on resource uptake (i.e., the amount of resource leaked into the environment depends on the amount being consumed). We use this specific structure as opposed to dependence of leakage directly on consumer biomass (e.g., [34], which would mean that the relative leakage to each resource type remains constant within a community), because it (i) more accurately reflects biological reality (microbes typically produce specific byproducts when feeding on specific resources [7]) and (ii) allows greater variation in cross feeding between consumer pairs (as the contribution to leakage from each consumer is unique) allowing us to better explore the effects of interactions. Also, we allow species to leak metabolites that they also consume (we address this assumption further in the Discussion).

We define the consumer’s maintenance cost to be:

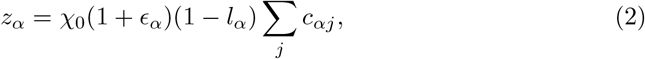

where χ_0_ is the average cost of being able to consume a given resource, the summation represents the total number of resources that species *α* is able to process, and *ϵ*_*α*_ is a small random fluctuation that introduces variation in the cost for species that have identical preferences. Eq 2 ensures that neither generalists nor specialists are systematically favoured during community assembly (by imposing a greater cost on species that consume a wider range of resources), and that all species are able to deplete resources to similar concentrations independently of their leakage level (see Supporting text section 1 for rationale, Fig S1 for results under different cost functions; and Discussion).

The above model entails the following assumptions: (i) all resources contain the same amount of energy (taken to be 1 for simplicity), (ii) a linear, non-saturating consumer functional response, (iii) binary consumer preferences (uptake rates), and (iv) an environment where all resources are externally supplied in equal amounts. We address the implications of these assumptions in the Discussion.

### Competition and facilitation metrics

In the system of equations 1, competition for resources exists because all pairs of consumer species generally share some resource preferences (their metabolic preferences vectors are not orthogonal). We quantify the pairwise competition between a species pair (*α, β*) by counting the resource preferences they share through the scalar product of their preference vectors, ***c***_*α*_ · ***c***_*β*_. Therefore, community-level competition (denoted as 𝒞) can be calculated by taking the average of the competition matrix, which encodes the competition strengths between all species pairs, that is

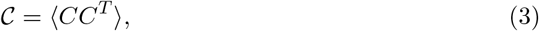

where *C* is the *s* × *m* matrix of metabolic preferences of all the species in the community.

On the other hand, facilitation occurs when a species leaks metabolic by-products that are used by another species. We measure pairwise facilitation from species *α* → *β* by calculating the fraction of secreted resources from species *α* that are consumed by species *β* per unit of resource abundance,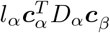. Similar to competition, we compute community-level cooperation (denoted as ℱ), by taking the average of the facilitation matrix, which encodes the competition strengths between all species pairs, that is

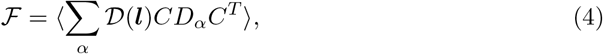

where where 𝒟(***l***) is a diagonal matrix with the leakage vector of each species in the community in its diagonal.

Henceforth, we refer to the quantity 𝒞 − ℱ as “net competition”, which we later show is related to the “cohesion” defined in previous work [25].

### Simulations

In Fig 1 we present an overview of our simulations, which we now describe. For the parameter values used, see Table 1.

**Table 1.**
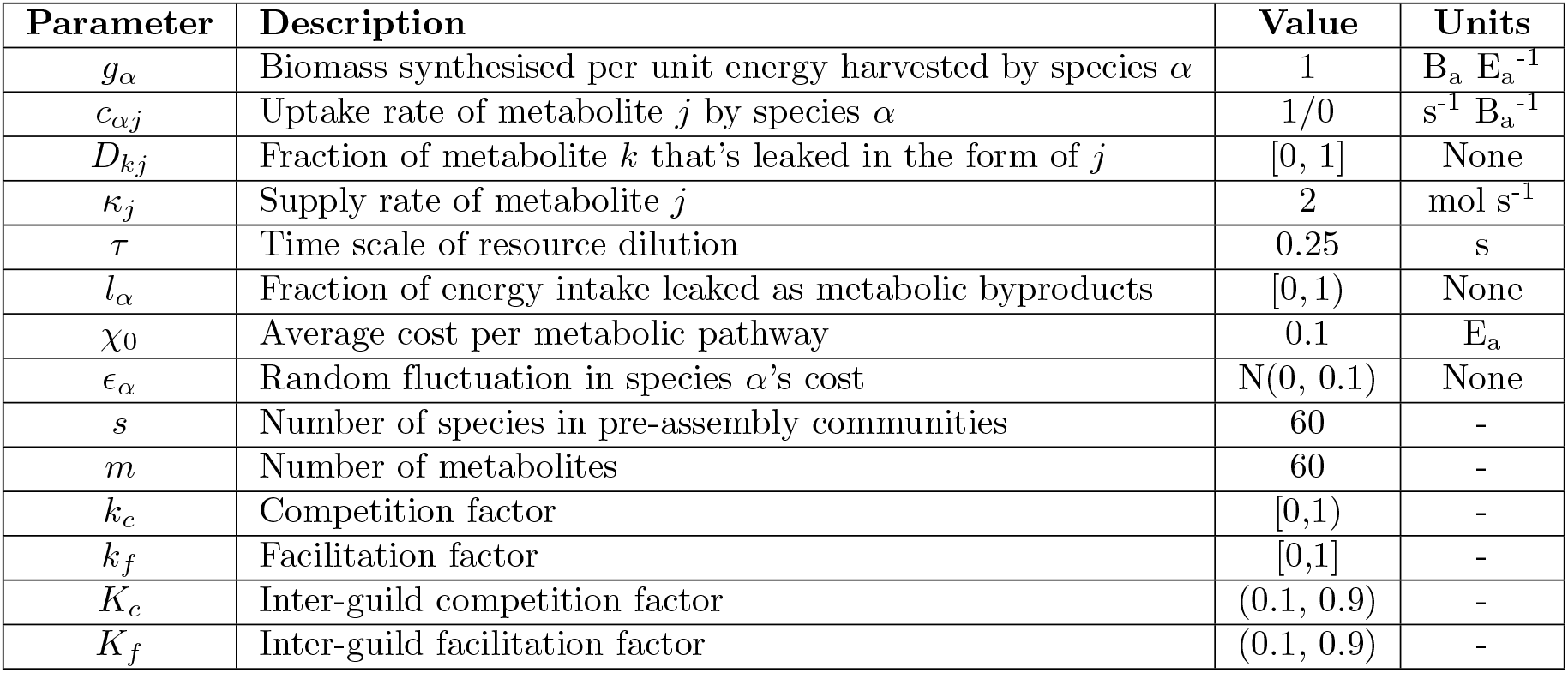
Table of parameters used in our model. The units E_a_ and B_a_ represent arbitrary energy and biomass units, respectively.

#### Step 1: Parameterization

We first set the parameters of the initial communities (before assembly) such that they span interactions across the spectrum of net competition (𝒞 − ℱ). For each parent community, we modulate the structure of the *C* and *D* matrices (consisting of the resource preferences *c*_*αj*_’s and secretion proportions *D*_*jk*_’s, respectively) by developing constrained random sampling procedures that guarantee specific levels of competition and facilitation at the community’s steady state (see Supporting text section 2). In addition, we also add structure to *C* and *D* to emulate the existence of distinct resource classes and consumer guilds (see Supporting text section 4). With these procedures, net competition in an assembled parent community can be regulated though four parameters: *k*_*c*_ (competition factor), *k*_*f*_ (facilitation factor), *K*_*c*_ (inter-guild competition factor), and *K*_*f*_ (inter-guild facilitation factor) (see Supporting text section 2). Note that we parameterize the initial communities by assuming (i) a shared core metabolism encoded in *D*, and (ii) a common leakage fraction *l* for all species (the implications of which we address in the Discussion), but we relax these assumptions in our coalescence simulations (Methods; Step 3).

#### Step 2: Assembly of parent communities

After parameterization, we numerically integrate Eqs 1 until steady state (a putative equilibrium) is reached. We perform 100 such assembly simulations with random sets of consumers for each combination of competition and facilitation factors (i.e., *k*_*c*_ = *k*_*f*_ ∈ [0, 0.5, 0.9]), repeating this for three values of leakage (*l* ∈ [0.1, 0.5, 0.9]). We compare species composition, abundances, and interaction structure of communities before and after assembly. In order to compare species composition, we calculate the difference between the proportion of species with a certain number of metabolic preferences, *n*_*r*_ (n-preference consumers, a measure of generalism), before and after assembly as

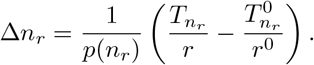

Here, *p*(*n*_*r*_) is the probability that a species has *n*_*r*_ metabolic preferences (Supporting text section 2), the 0 denotes before assembly, *r* is species richness, and *T*_*n*_*r* is the number of species with *n*_*r*_ preferences. Thus, when Δ*n*_*r*_ > 0 the proportion of species with *n*_*r*_ metabolic preferences increases after assembly and vice versa. In order to analyze species abundances, we track the abundance fraction of consumers in each group of n-preference species, calculated as total abundance of the species in the group, divided by the total community biomass. Finally, we address the interaction structure after assembly by quantifying the levels of competition (𝒟) and facilitation (ℱ) in the assembled communities (Fig 2A).

**Fig 2.**
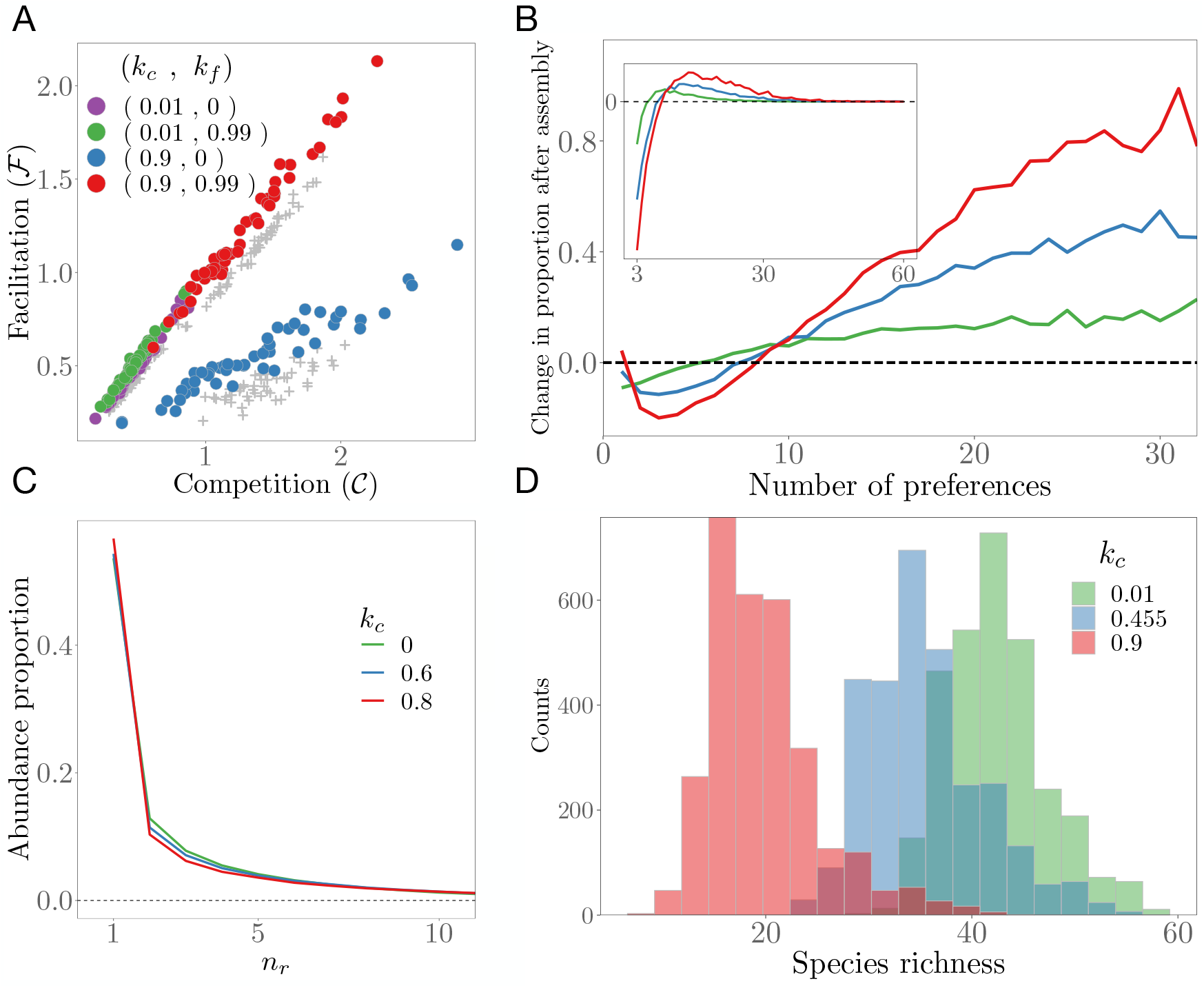
Features of assembled parent communities. **A**: Facilitation versus competition level in starting (grey dots) and assembled (coloured dots) communities for leakage *l* = 0.9 and different combinations of competition (*k*_*c*_) and facilitation (*k*_*f*_) factors. Communities assembled for each pair of [*k*_*c*_, *k*_*f*_] values have the same colour. The assembled communities are always significantly more cooperative at the end of the assembly than at the start. **B**: Change in proportion of species in each n-preference consumer group (Δ*n*_*r*_, Methods; Step 2) before and after assembly, for different values of *k*_*c*_ (legend in panel C) indicating that more generalist species are less prone to extinction during assembly. Values for *n*_*r*_ > 30 had too much uncertainty due to low sampling and therefore have been removed for clarity. **Inset:** Δ*n*_*r*_ is weighted by the abundance fraction at equilibrium of each n-preference consumers group. **C**: Abundance fraction of the n-preference species groups for different values of *k*_*c*_. **D**: Distributions of species richness values of parent communities assembled under different *k*_*c*_ values. Increasing competitiveness tends to decrease species richness.

#### Step 3: Coalescence

To simulate coalescence between a pair of assembled parent communities, we set all resources to their initial concentrations, and numerically integrate the new combined system to steady state. In order to disentangle the effects of competition versus cooperation and study the effect of repeated coalescence events, we simulate three coalescence scenarios: random, recursive, and serial, as follows (further details in Supporting text section 3).

##### Random coalescence

To address the effects of competition alone in the outcome of coalescence events, here we coalesce pairs of randomly sampled parent communities having the same leakage value *l* (2 ·10^4^ pairs for each leakage level, Fig 3C). That is, we fix the leakage level to ensure that the communities have, on average, similar cooperation levels, but leave *k*_*c*_ free to vary such that they span a broad range of competition levels.

**Fig 3.**
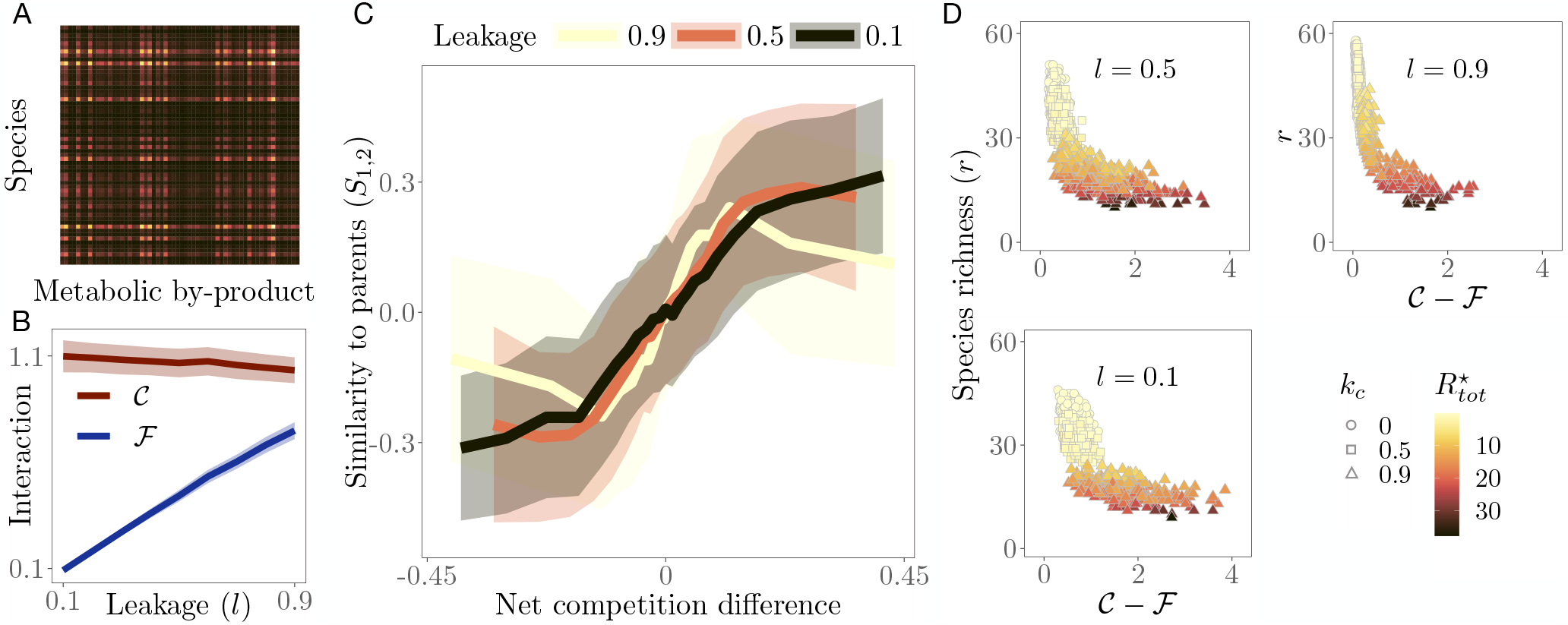
Community coalescence between pairs of randomly picked communities with same leakage. **A**: Example of the secretion matrix with elements (*CD*)_*αk*_ representing the total leakage of resource *k* by species *α*. **B**: Community-level competition 𝒞 (dark red) and facilitation ℱ (blue) averaged across simulations for each leakage value. Since competition does not depend on the leakage, it remains consistently high throughout. Facilitation, on the other hand, increases linearly with leakage. **C**: Parent community dominance (*S*_1,2_) as function of net competition difference (𝒞 _1_− ℱ_1_) (𝒞_2_− ℱ_2_) (solid lines ± 1 standard deviation (shaded)), binned (20 bins) over communities with similar x axis values, for three community-wide leakage levels. The post-coalescence community is more similar to its less (net) competitive parent. **D**: Species richness (*r*) as a function of net competition in parent communities, coloured by total resource concentration at steady state 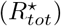. The observed negative correlation for all values of leakage shows that communities with lower net competition tend to be more species-rich and also better at depleting resources (brighter coloured values, corresponding to lower levels of 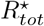 are scattered towards the top left of the plots).

##### Recursive coalescence

In order to study the effects of cooperation in particular on community coalescence, we repeatedly coalesce a given pair of communities *A* and *B*, slightly increasing the leakage of the latter in each iteration (Fig 4A). This allows us to modify the strength of cooperative interactions in the community, because facilitation is proportional to *l* (Eq 4), while keeping competition levels constant, because competition is independent of *l* (Eq 3) and the remaining parameters are kept fixed.

**Fig 4.**
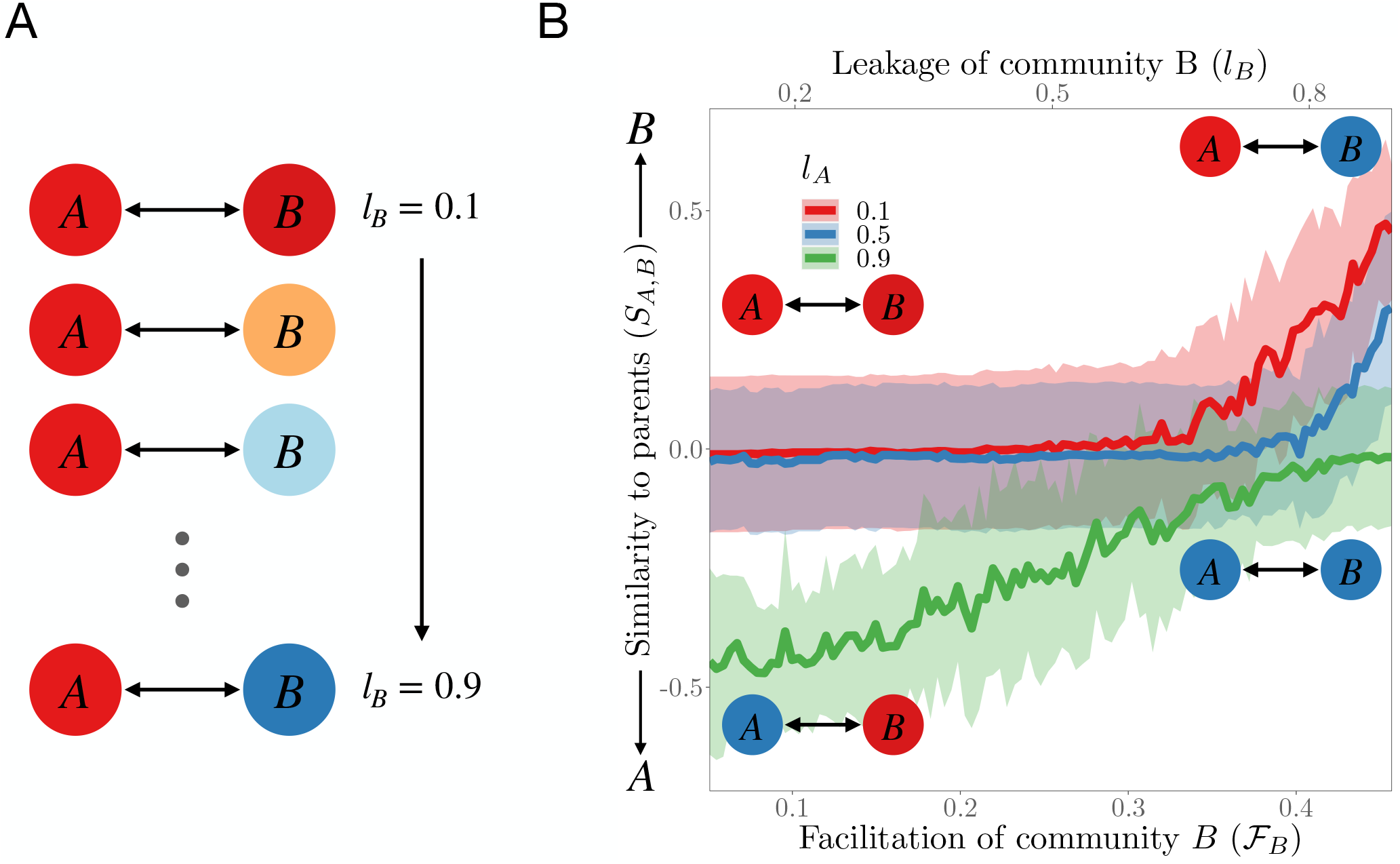
Recursive coalescence between microbial communities. **A**: Sketch of the simulation set up. The same pair of communities (*A, B*) is recursively coalesced, with the leakage of *B* gradually increasing after each coalescence event for three levels of leakage of A, and 25 replicates per *l*_*A*_ value. **B**: Parent community dominance after coalescence between communities *A* and *B*, as a function of facilitation level of community *B, ℱ* _*B*_ (bottom x axis), and leakage of community *B, l*_*B*_ (top x axis). Each curve corresponds to a different value of *l*_*A*_. Shaded regions are ±*σ*. Dominance of parent community *B* after coalescence increases with *l*_*B*_, implying that higher cooperation levels enhance coalescence success.

##### Serial coalescence

In the natural world, a community may be exposed to more than one coalescence event. Consequently, here we simulate a scenario where a local (“resident”) community ℛ harbouring species with leakage *l*_ℛ_, and metabolism *D*_ℛ_ is successively invaded by many other randomly sampled communities (“invaders”), ℐ with species of leakage *l*_ℐ_ and metabolism *D*_ℐ_ (Fig 5A). This allows us understand how the functional and structural properties of a microbial community evolve over time under successive encounters with other communities.

**Fig 5.**
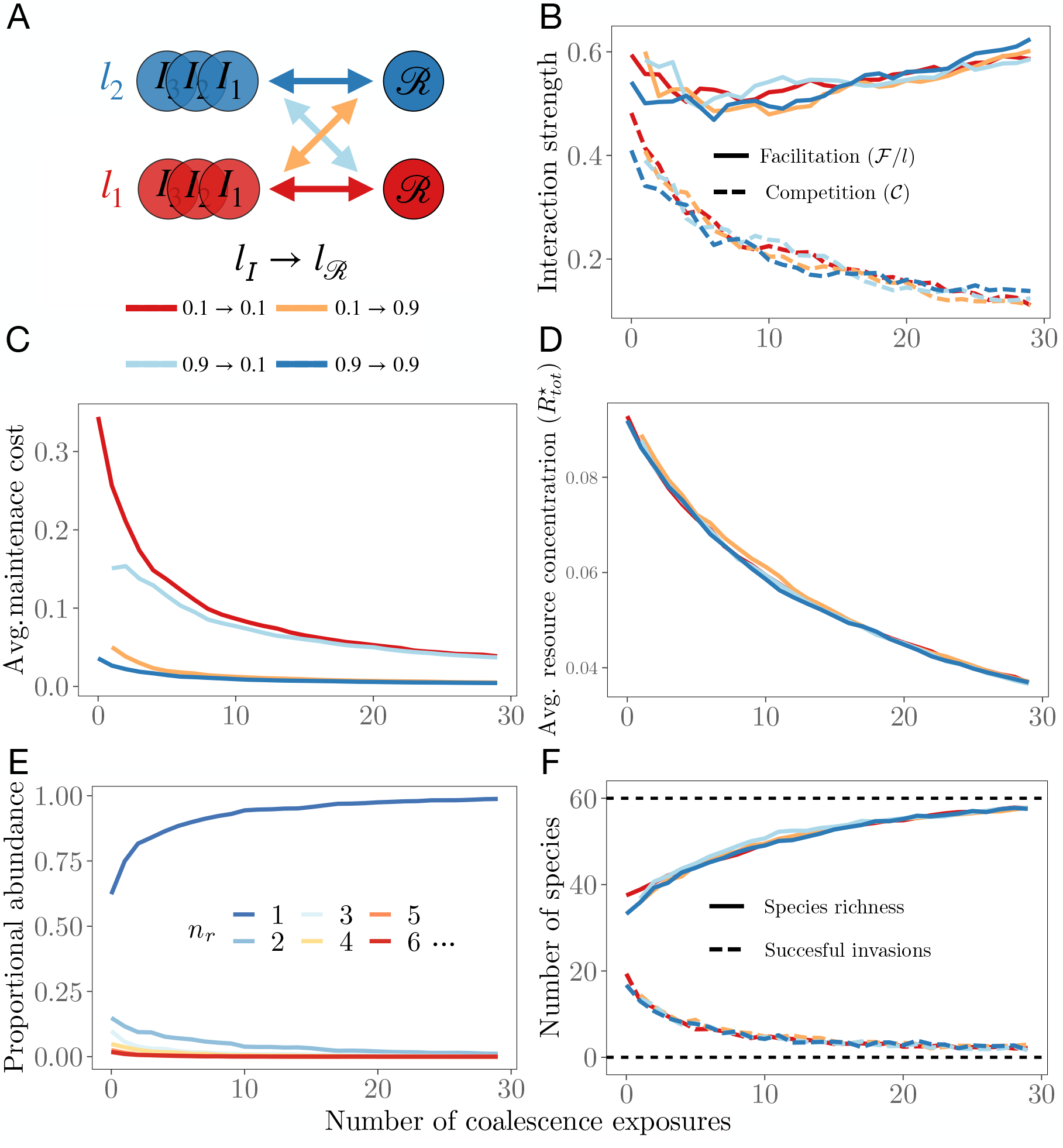
Serial coalescence of microbial communities. **A**: Sketch of the simulation set up. Resident communities (ℛ, upper circles) with leakage *l*_ℛ_ are successively coalesced with randomly sampled invader communities (ℐ, lower circles) with leakage *l*_ℐ_ for all possible combinations of leakage values (arrows) ***l***_ℐ_ = ***l***_ℛ_ = [0.1, 0.9]. For each serial coalescence sequence, we examine as a function of number of coalescence events, the following community properties of ℛ: (**B**) community-level competition (𝒞, dashed lines) and facilitation (ℱ, solid lines); (**C**) average species maintenance cost; (**D**) average resource concentration at equilibrium; (**E**) abundance fraction of each n-preference species group; and (**F**) number of successful invasions and with species richness. All the measures are averaged across 20 replicates. The standard deviation is decreasing along the x-axis and is never more than 40% of the mean for all the curves (not shown to reduce clutter). Abundance fraction of species with *n*_*r*_ > 5 was negligible and is not plotted for clarity.

At the end of each random and recursive coalescence simulation, we quantify the dominance of either parent community in the post-coalescence community by measuring similarity of the latter to each of the two parents (indexed by 1 and 2) as:

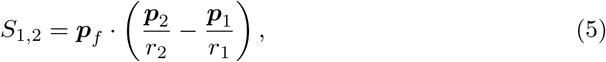

where ***p***_*f*_, ***p***_1_, and ***p***_2_ are (*s*_1_ + *s*_2_)–dimensional vectors of species presence-absence in the post-coalescent, and parent communities 1, and 2, respectively, with *r*_1_ and *r*_2_ the species richness values of the parent communities 1 and 2, respectively (calculated as *r*_*i*_ = Σ*p*_*i*_). If *S*_1,2_ = −1, the coalesced community is identical to parent community 1, and if *S*_1,2_ = 1, it is identical to parent community 2. This measure is independent of the species richness. Thus we can mix communities with different species richness while avoiding a bias in similarity towards the richer one. We then analyze how this dominance measure depends on the interaction structure of the parent communities (𝒞 _1,2_ and ℱ_1,2_; Eqs 3 and 4). After each coalescence event in the serial coalescence procedure, we measure competition and facilitation levels of the resident community, along with the average species maintenance cost, average resource abundance at equilibrium, species richness, and number of successful invasions, during the entire sequence of serial coalescence events. For all assembled parent as well as coalesced communities we confirmed that the steady state was a locally asymptotically stable equilibrium point (Supporting text section 1).

## Results

### Assembly of parent communities

In Fig 2 we show the key features of assembled communities. Figure 2A shows that as expected from Eqs 3 and 4, the levels of community-wide competition and facilitation are positively correlated, mediated by the structure of the *C* and *D* matrices. Figure 2B shows that the difference between the proportion of n-preference consumers before and after assembly (Δ*n*_*r*_, Methods; Step 1), increases for all simulated values of *k*_*c*_, indicating that more generalist species are less prone to extinction during assembly. For the lowest value of *k*_*c*_, Δ*n*_*r*_ is in fact a monotonically increasing function of the number of preferences. This is expected because a species able to harvest energy from multiple resource pools is less likely to go extinct during community dynamics. As *k*_*c*_ increases, Δ*n*_*r*_ reaches a minimum (Fig 2B), indicating that in more competitive environments pure specialists become more prevalent than moderate generalists. This is due to the fact that in a highly competitive environment the resource demands are concentrated on a subset of resources, while others are barely consumed (Supporting text section 2). In these communities, consumers that specialize exclusively on empty niches thrive.

Figure 2C shows that specialist consumers are systematically present in higher abundance than generalists for all values of *k*_*c*_. This is because several specialists are able to deplete all resources through their combined action more efficiently than one generalist [26], and as a result, although generalists are more persistent than specialists upon assembly (Fig 2B), they achieve lower abundances at equilibrium (Fig 2C). In Figure 2B (inset), Δ*n*_*r*_ is weighted by the abundance fraction (Methods; Step 2) of each group of n-preference consumers. This reveals an optimal group of consumers with a number of metabolic preferences that maximizes both survival probability and abundance at equilibrium. This optimal value increases for more competitive environments (as *k*_*c*_ increases). Finally, Figure 2D shows that more competitive communities tend to be less species rich, as expected from general competition theory.

### Reducing competition increases coalescence success

Figure 3 shows that communities with lower net competition values tend to perform better in coalescence as seen by the positive relationship between parent community dominance (*S*_1,2_) and the quantity (𝒞_1_ − ℱ_1_) − (𝒞 _2_ − ℱ_2_) (Fig 3C). That is, communities that emerge following coalescence tend to have greater similarity with the less net competitive parent. This trend holds at higher values of leakage, where cooperation levels are significant (Fig 3B), but with a clear critical point (the yellow line reverses in direction at a value of effective competition difference). This pattern is driven by the fact that less competitive parent communities deplete resources more efficiently and achieve a higher species richness (Fig 3D; Supporting text section 1). All these results also qualitatively hold for microbial communities that have consumer guild structure (Supporting text section 4).

### Cooperation further enhances coalescence success

Figure 4 shows that when a community (*B*) whose leakage fraction increases successively during recursive coalescence events with another (*A*) with a fixed leakage level, the former becomes increasingly dominant. The result is consistent for a range of leakage values of community *A*. This shows that increasing cooperation levels enhance coalescence success.

### Community evolution under repeated coalescence events

Figure 5 shows that on average, competition level significantly reduces and facilitation level increases during repeated coalescence events. Along with this, the average maintenance cost of species present in the resident community decreases with the number of coalescence exposures, and so does average resource abundance at equilibrium (Fig 5C and D), indicating that resource depletion ability improves in the process. In addition, the sub-population of resource specialists (that consume only one resource) increases with the number of coalescence events, while the rest of the species groups decrease in abundance (Fig 5E). Finally, the number of successful invasions into the resident community decreases function of number of coalescence events, while its species richness increases (Fig 5F). Taken together, these results show that communities composed of non-competing specialists that cooperate among themselves (Fig 5B and E) and reduce their respective metabolic costs (Fig 5C), improve their overall resource depletion ability (Fig 5D). This, in turn, makes them more resistant to multi-species invasions and therefore more successful in pairwise coalescence events (Fig 5F).

## Discussion

Our findings offer new mechanistic insights in the dynamics and outcomes of microbial community coalescence by explicitly considering the balance between competition and cooperation; two key interactions of real microbial communities [26, 37]. Specifically, we find that communities harbouring less competing and more cooperative species (that is, having lesser net competition) dominate after coalescence because they are better at depleting resources and resisting invasions. Therefore, when a community undergoes a series of coalescence events, its competitiveness decreases and cooperativeness increases, along with its species richness, resource use efficiency, and resistance to invasions. These results provide a theoretical foundation for hypotheses suggested recently [18, 21], and mechanistic insights into empirical studies that have demonstrated the importance of cross-feeding interactions on community coalescence [22].

Our result based on coalescence between pairs of random communities at very low leakage (black line in Fig 3C) essentially extends the results of [25] to communities with both competitive and cooperative interactions. Tikhonov showed that coalescence success is predicted by minimizing community-level competition through the optimisation of resource niche partitioning, which also guarantees maximization of resource depletion efficacy. Here we show that, similarly, the successful community is the one that achieves lower *net* competition (𝒞 − ℱ), which also predicts community-level resource depletion efficacy as well as species richness (Fig 3D). Thus, simultaneously reducing competition and increasing cooperation together drives the outcome of community coalescence. Therefore, the quantity – (𝒞 − ℱ) = ℱ − 𝒞 is also a measure of the “cohesiveness” of a microbial community. However, we also find that at extreme value of leakage (*l* = 0.9), there is a critical level of net competition difference beyond which coalescence success decreases again (yellow line in Fig 3C). This suggests that in the regime of high cooperation and competition, (high leakage, and tail ends of the curve) facilitative links in fact become detrimental. A similar result has been reported reported in [65]. This critical value is not seen when the cost function does not include leakage (Fig. S2). Interestingly, we also find that this phenomenon is very weak when biologically-realistic guild structure is present (Fig. S6). These effects of extreme leakage (and facilitation) on coalescence success cannot be predicted by our model analyses (Supporting text section 1), and merit further investigation in future research, provided such high leakage levels are biologically feasible.

In our model systems, species compete not only for resources leaked by other species, but also for resources leaked by themselves, i.e., species may leak metabolic by-products that are also encoded in their consumer preferences vector. Leakage of metabolic resources is a pervasive phenomenon in the microbial world [38, 39], and has been shown to exist also in resources necessary for growth, even in situations when those essential metabolites are scarce [40, 41]. Although it may seem counter-intuitive for microbes to secrete metabolites essential for their own growth, such leakage can be advantageous, especially in bacteria, as “flux control” or growth-dilution mechanisms which provide short-term growth benefits in crowded environments [42, 43].

Our recursive coalescence simulations (Fig 4A) allowed us to establish that coalescence success is enhanced by cooperative interactions. This result is consistent with past theoretical work showing that mutualistic interactions are expected to increase structural stability by decreasing effective competition [44]. It is also consistent with recent theoretical results on single species invasions in microbial communities [45]. Nonetheless, this finding hinges on our choice of the cost function (Eq (2); Supporting text section 1). This cost function, which was motivated by biological considerations, imposed an efficiency cost to species with lower leakage, ensuring that all consumers, independently of their leakage fraction, depleted resources to the same concentration on average (see Supporting text section 1). This allowed us to perform coalescence events between communities harbouring species with different leakage without introducing a bias towards the more efficient species. This choice corresponds biologically to the interpretation of leakage as an efficiency factor in the conversion from energy to biomass (Eq S2 in Supplementary information). As a consequence, higher leakage species reach, in general, lower abundances at equilibrium [46].

Our findings about the evolution of community-level properties in response to repeated community-community encounters (Fig 5) suggest that it might be possible to identify functional groups of microbes or microbial traits that are are a “smoking gun” of past coalescence events experienced by a given community [18]. Additionally, our finding that members of communities with a history of coalescence are likely to be become increasingly resistant to further community invasions suggests a novel and potentially economical way to assemble robust microbial communities. We also found that repeated coalescence events contributed to increase species richness, offering another mechanism that may help explain differences in microbial diversity across locations and environments [18].

Our finding that resident communities exposed to repeated community invasions were mainly composed of cooperative specialists (Fig 5B and E) is due to our assumption that all resources were supplied, and at a fixed rate. This allowed specialists to survive because their only source of energy was always provided. This property may not be as commonly seen in real communities, where fluctuations in resource supply are common. Ignoring environmental fluctuations allowed us to focus on coalescence outcomes in terms of the species interaction structure alone. While this assumption may be sensible in some cases [47], it is an oversimplification in others [33]. Therefore, studying the complex interplay between biotic interactions and environmental factors, e.g., by allowing substrate diversification from a single supplied resource [6, 7], or perturbing the supply vector periodically to simulate some form of seasonality, is a promising direction for future research. In such cases, we expect a more balanced mix of generalists and specialists, such that only the competitive interactions necessary to diversify the available carbon sources will persist upon coalescence events, but above that threshold, the results presented here (Figs 3, 4, and 5) would be recovered.

Assuming a core leakage and metabolism common to the whole community, made the assembly dynamics computationally tractable, while ensuring that the system was not far away from the conditions of real communities [6, 27]. The assumption of common leakage was relaxed in the recursive coalescence procedure. In addition, both the community-wide fixed leakage and core metabolism assumptions were relaxed in the serial coalescence simulations, thus using the model in its fully general form as seen in Eqs 1. Finally, to retain an analytically tractable theoretical setting for an otherwise complex system, we assumed binary consumer preferences, linear consumer functional responses, and resources of equal energetic value throughout. While this is a promising avenue for future work, we expect our results to be qualitatively robust to relaxation of these constraints, based on recent work on microbial community assembly dynamics using the same general model [6, 27].

Encounters between microbial communities are becoming increasingly frequent [52], and mixing of whole microbial communities is gaining popularity for bio-engineering [53], soil restoration [54], faecal microbiota transplantation [55, 56], and the use of probiotics [57]. We present a framework which relates the structure of species interactions in microbial communities to the outcome of community coalescence events. Although more work is required to bridge the gap between theory and empirical observations, this study constitutes a key step in that direction.

## Supporting information

**Supporting text section 1 – Further details of the mathematical model**.

**Supporting text section 2 – Modulating net competition levels**.

**Supporting text section 3 – Further details of the community coalescence simulations**.

**Supporting text section 4 – Adding consumer guild structure**.

**Fig S1** Consequence of relaxing the assumption of leakage dependence.

**Fig S2** Results of random coalescence procedure without imposing the efficiency cost (leakage dependence).

**Fig S3** Eigenvalues of J when evaluated at equilibrium for two example communities.

**Fig S4** Examples of differently-structured preference (*C*) and metabolic (*D*) matrices.

**Fig S5** Auto-correlation of vector of species abundances in community B for consecutive re-assemblies of this community, along the studied leakage range.

**Fig S6** Community coalescence with consumer guilds present.

## Author Contributions

**Conceptualization:** Pablo Lechón, Tom Clegg, Jacob Cook, Thomas P. Smith, Samraat Pawar

**Methodology:** Pablo Lechón

**Software and Visualizations:** Pablo Lechón

**Writing – Original Draft Preparation:** Pablo Lechón

**Writing – Review & Editing:** Pablo Lechón, Tom Clegg, Jacob Cook, Thomas P. Smith, Samraat Pawar.

**Funding Acquisition:** Samraat Pawar

## Notes

### Competing Interest Statement

The authors have declared no competing interest.

### Summary of Updates

Submitted version

