## Supplementary material for "The role of competition versus cooperation in microbial community coalescence"

September 1, 2021

### S1 Further details of the mathematical model

The model is (same as main text Eq 1):

$$\begin{aligned} \frac{dN_\alpha}{dt} &= g_\alpha N_\alpha \left( (1 - l_\alpha) \sum_j c_{\alpha j} R_j - z_\alpha \right), \\ \frac{dR_j}{dt} &= \kappa_j + \tau^{-1} R_j - \sum_\alpha N_\alpha c_{\alpha j} R_j + \sum_{\alpha k} N_\alpha l_\alpha D_{\alpha k j} c_{\alpha k} R_k. \end{aligned} \tag{S1}$$

In matrix form, this system can be written as as:

$$\begin{aligned} \frac{d\boldsymbol{N}}{dt} &= \mathcal{D}(\boldsymbol{g} \circ \boldsymbol{N}) \left( \mathcal{D}(\mathbb{1} - \boldsymbol{l}) C \boldsymbol{R} - \boldsymbol{z} \right) \\ \frac{d\boldsymbol{R}}{dt} &= \boldsymbol{\kappa} - \tau^{-1} \boldsymbol{R} - \mathcal{D}(\boldsymbol{R}) C^T \boldsymbol{N} + \sum_\alpha l_\alpha D_\alpha^T \mathcal{D}(\boldsymbol{R}) \boldsymbol{c}_\alpha N_\alpha \end{aligned}$$

Here, $\circ$ denotes an element-wise operation, $\boldsymbol{R}$ and $\boldsymbol{N}$ are the vectors of resource concentrations and population abundances respectively, $\mathbb{1}$ is a vector of ones, and the remaining symbols in boldface represent the same quantities in Eq S1 in vector form. $C$ is the matrix of consumers' preferences, and $\mathcal{D}(\boldsymbol{x})$ is a diagonal matrix with the vector $\boldsymbol{x}$ as its diagonal.

#### S1.1 Cost function

Every microbial cell has a maintenance cost, which is the energy required to perform tasks to remain alive such as transportation of metabolites and synthesis of RNA and enzymes for metabolizing substrates. We define this cost to be

$$z_\alpha = \chi_0 (1 + \epsilon_\alpha)(1 - l_\alpha) \sum_j c_{\alpha j}.$$

Here, $\chi_0$ is the average cost of consuming a given resource, the summation represents the total number of resources that species $\alpha$ is able to process, and $\epsilon_\alpha$ is a small random fluctuation sampled from a truncated normal distribution ($\epsilon_\alpha \sim N(0, 0.1)$), that introduces variation in the cost for species that have identical preferences. Also, note that the fitness of an organism is determined by both its cost and its metabolic preferences, so we keep the random fluctuations to be small values (relative to the uptake values) in all simulations because if the are too large, fitness of the organism would be purely determined (randomly) by its cost, which would be biologically unrealistic considering the unavoidable feedback between environment and consumers in determining their fitness. As such, setting $\epsilon_\alpha = 0 \; \forall \; \alpha$ would not qualitatively change our results about coalescence outcomes.

This cost function entails two key assumptions. The first assumption is that generalist consumer species (which feed on a wide range of resources types) pay a higher maintenance cost (the summed term in the cost function) than specialists (which consume one or few resource types; henceforth, the "generalism cost"). More generalist species necessarily maintain more complex metabolic networks than specialist species, and the upkeep of larger metabolic networks (and thus, larger genomes) incurs in greater maintenance costs (DeLong et al. 2010,

Kempes et al. 2017), leading to a trade-off between resource generalism and cost of maintenance. Here we impose this trade-off such that the maintenance cost is proportional to the sum of all resource preferences (i.e. the number of resources consumed). This assumption is similar to the one made by Tikhonov (2016) (also see Tikhonov & Monasson (2017)).

The second assumption is that the cost of cellular maintenance is also proportional to the fraction of retained resources $(1 - l)$, i.e., species that retain greater quantities of resources (those with less leakage, $l$) have higher maintenance costs (the "efficiency cost"). This assumption stems from the fact that the processing of resources itself incurs a metabolic cost (WIESER 1994). Therefore those species retaining greater quantities (leaking less) of resources must pay a greater metabolic cost of processing those resources. With this cost function, the consumer equation from Eq S1 becomes

$$\frac{dN_\alpha}{dt} = g_\alpha N_\alpha (1 - l_\alpha) \left( \sum_j c_{\alpha j} R_j - \chi_0 (1 + \epsilon_\alpha) \sum_j c_{\alpha j} \right). \tag{S2}$$

Thus, leakage is now an factor that also determines what proportion of the harvested energy is be converted to biomass, consistent with the classical notion of biomass conversion or assimilation efficiency (WIESER 1994).

#### S1.2 Effect of the cost function on community feasibility and dominance

We now explain how the above maintenance cost function has important implications for community feasibility and ultimately, coalescence outcomes. Species $\alpha$ will reach a feasible (positive) equilibrium when its resource surplus term, that is, the amount of energy left when the maintenance cost is subtracted from the initial harvest (terms inside the brackets of the consumer equation in Eq S1) equals 0. That is (after substituting the cost function),

$$\frac{1}{g_\alpha N_\alpha} \frac{dN_\alpha}{dt} = (1 - l_\alpha) \left( \sum_j c_{\alpha j} R_j - \chi_0 (1 + \epsilon_\alpha) \sum_j c_{\alpha j} \right) = 0.$$

Therefore, at steady state,

$$\sum_j c_{\alpha j} R_j^* = \chi_0 (1 + \epsilon_\alpha) \sum_j c_{\alpha j}. \tag{S3}$$

For the whole community, the vector of equilibrium resource abundances is given by,

$$\begin{aligned} C\boldsymbol{R}^\star &= \chi_0 \mathcal{D}(\mathbb{1} + \boldsymbol{\epsilon}) C \mathbb{1}, \\ \text{i.e., } \boldsymbol{R}^\star &= \chi_0 C^{-1} \mathcal{D}(\mathbb{1} + \boldsymbol{\epsilon}) C \mathbb{1} \end{aligned} \tag{S4}$$

Here, $\boldsymbol{R}^\star$ is the vector of resource concentrations (the $^\star$ indicating it's equilibrium state), $C$ is the consumer preferences matrix, $\mathbb{1}$ is a vector of ones, and $\mathcal{D}(\mathbb{1} + \boldsymbol{\epsilon})$ is a diagonal matrix with the vector $\mathbb{1} + \boldsymbol{\epsilon}$ (the vector of random fluctuations in costs) in its diagonal. As such, Eq. (S3) (and (S4)) is a necessary (but not sufficient) condition for the community's feasibility (existence of the consumer-resource equilibrium) provided that (i) the preferences matrix $C$ is invertible, and (ii) all the resource concentrations at equilibrium are positive. In the following analysis, we only consider systems where these two conditions are satisfied, because our goal is to establish the conditions for community dominance following coalescence *given* a pair of feasible (and locally asymptotically stable; next section) parent communities. Our simulations show that these results hold even when we coalesce communities with feasible equilibria consisting of different consumer numbers (the measure $S_{1,2}$ is independent of species richness; main text Methods, Step 3).

We now show that the two key assumptions of our cost function—the efficiency cost and the generalism cost—are not just biologically realistic but also crucial for understanding the role of competition and cooperation in community coalescence. To this end, we relax each assumption, and re-calculate the resource abundance vector at equilibrium. When we relax the assumption of an efficiency cost, the equilibrium resource vector becomes

$$\boldsymbol{R}^{\star} = \mathcal{D}^{-1}(\mathbb{1} - \boldsymbol{l})\chi_0 C^{-1}\mathcal{D}(\mathbb{1} + \boldsymbol{\epsilon})C\mathbb{1}. \tag{S5}$$

In this case, in contrast to Eq. (S4), the equilibrium resource abundances (i.e., magnitude of the abundance vector $\boldsymbol{R}^{\star}$) increases with values of species' leakages (magnitude of $\boldsymbol{l}$). Thus, all else (including the supply of different resources) being the same, a "less leaky" community will deplete the given resource pool to a lower concentration than a more leaky one. This means that when two such communities are coalesced, the one with lower leakage will create an environment that does not guarantee feasible coexistence of consumers in the other community (maintenance cost becomes higher than energy harvest, resulting in negative growth rates). That is, upon community-community encounter, more consumer species of the leakier community will be driven extinct relative to those from the less leaky one because the resource environment a not capable of sustaining a positive equilibrium. In other words, in the absence of an efficiency cost, communities with higher leakage will be systematically be displaced or dominated by communities with lower leakage. This can be interpreted as a variant of Tilman's $R^{\star}$ rule applied to a pair of communities instead of a pair of species (also see Tikhonov (2016)). This analytical result is demonstrated numerically in Fig S1. Furthermore, since leakage is directly correlated with community-wide cooperation level (main text Eq. 4), relaxing this assumption would imply that being more cooperative is detrimental for coalescence success, contradicting current literature which has found that cooperation generally increases structural stability and resistance to invasions (Pascual-García & Bastolla 2017, Kurkjian et al. 2021).

Next, we eliminate the generalism cost, which means that the equilibrium resource vector becomes

$$\boldsymbol{R}^{\star} = \chi_0 C^{-1}(\mathbb{1} + \boldsymbol{\epsilon}).$$

In this case, magnitude of the resource abundance vector at equilibrium decreases with the number of consumer species' resource preferences (magnitude of $C$). This means that generalist consumers are able to deplete resources more efficiently than specialists just because they posses more metabolic pathways, without incurring in an extra cost (i.e., a species able to consume 5 resources would harvest 5 times more energy than a species with just 1 metabolic preference). However, this is unrealistic, since, as mentioned above, the more metabolic pathways present, the higher the probability that any two of them require different

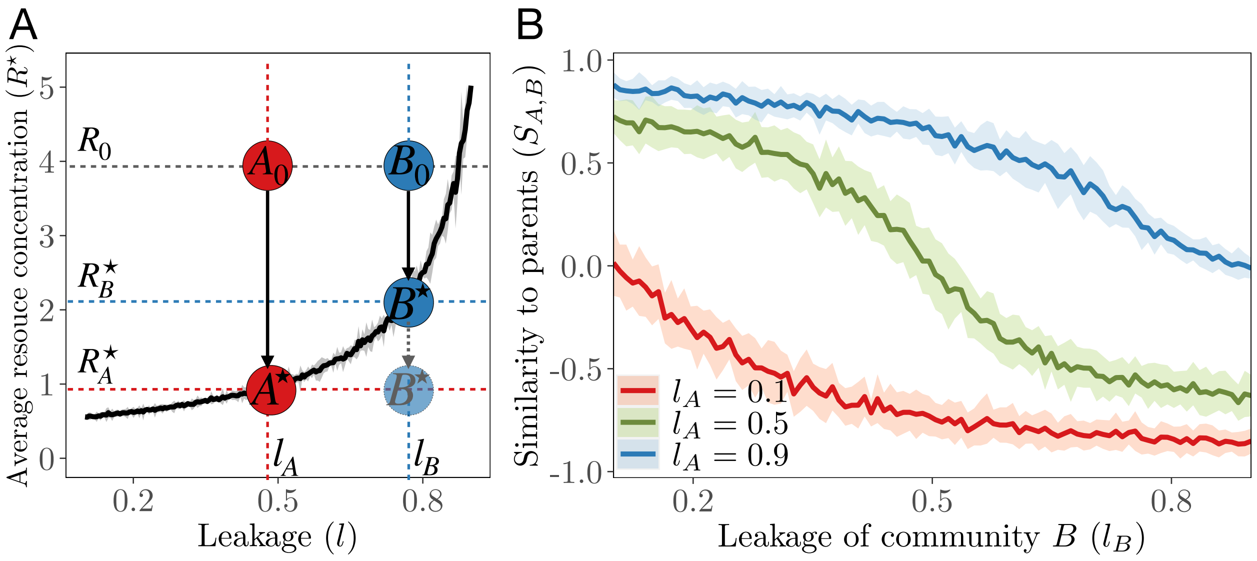

Figure S1: **Consequence of eliminating leakage-dependence of the cost function for community dominance in pairwise coalescence events.** **A**: Without leakage dependence of the cost function, mean of its resource concentrations ($R^\star$) at equilibrium (black curve) increases with leakage. The black curve is the mean of the mean resource concentrations (shaded region is $\pm\sigma$) reached at equilibrium for many random instances of feasible parent communities. Therefore, when any two communities with different leakage values $A$ ($l_\alpha \; \forall \; \alpha = l_A$) and $B$ ($l_\alpha \; \forall \; \alpha = l_B$) (all species set to have the same value for simplicity) are assembled in isolation in the same environment $R_0$, they will deplete the resources to concentrations $R_A^\star$ and $R_B^\star$, respectively. As a result, when the two are coalesced, community $A$, which can deplete the resources to a lower concentration, will create an environment that does not guarantee feasibility of consumers in community $B$ (semi-transparent blue circle), causing more species from community B to be driven extinct relative to those from A. **B**: Parent community dominance ($S_{A,B}$) after repeated coalescence events between pairs of communities of the types $A$ and $B$. We use the recursive coalescence simulation procedure (see main text Methods, Step 2 and Section S3), where the leakage ($l_B$, x-axis) is slightly increased after each iteration. So this simulation is equivalent to that producing main text Fig 4B, the only difference being that here the cost function lacks the leakage term. The result here is opposite to the one in main text Fig 4B, confirming that if the cost function is independent of leakage level, lower leakage (and therefore less cooperation) favors parent community dominance after coalescence. Note the sharper decrease in community dominance of $B$ as its increasing leakage ($l_B$) approaches the leakage of its competitor ($l_A$).

cellular machinery to be activated for optimal maintenance and functioning (e.g., through two different cellular compartments), incurring extra cost (Tikhonov & Monasson 2017).

Therefore, both efficiency and generalism costs are necessary for a meaningful analysis of the effect of cooperation vs competition on community coalescence outcomes. Of course, if two communities have the same leakage levels, they will on average have similar levels of cooperation, in which case competition level is the only factor that needs to be considered, and the presence of an efficiency cost does not matter. To confirm this, we ran the random coalescence procedure with communities assembled with a cost function lacking leakage dependence (Fig S2), which, as expected, yields qualitatively the same result as for communities assembled *with* the efficiency cost (main text Fig 3) because we are only coalescing communities with the same leakage value.

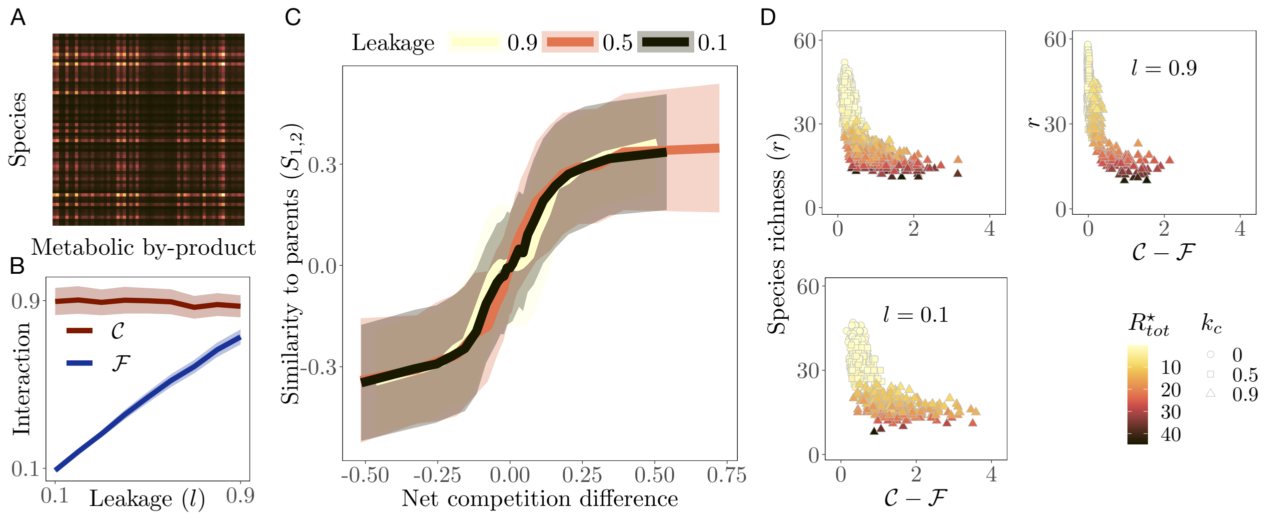

Figure S2: **Results of random coalescence procedure without imposing the efficiency cost (leakage dependence).** See main text Fig 4 for descriptions of the sub-panels.

#### S1.3 Stability

After assembly of a feasible parent or coalesced community, we assessed its local stability as follows. We first compute the Jacobian matrix of the system, and evaluate it at the steady state population and resource abundances (the feasible equilibrium, which may or may not be stable). For our system, the Jacobian is the block matrix of the form

$$
\begin{aligned}
J\Big|_{\substack{\boldsymbol{N}=\boldsymbol{N}^\star\\ \boldsymbol{R}=\boldsymbol{R}^\star}} &= \begin{bmatrix} \dfrac{\partial \dot{\boldsymbol{n}}}{\partial \boldsymbol{n}} & \dfrac{\partial \dot{\boldsymbol{n}}}{\partial \boldsymbol{R}} \\[2ex] \dfrac{\partial \dot{\boldsymbol{R}}}{\partial \boldsymbol{n}} & \dfrac{\partial \dot{\boldsymbol{R}}}{\partial \boldsymbol{R}} \end{bmatrix}\Bigg|_{\substack{\boldsymbol{n}=\boldsymbol{N}^\star\\ \boldsymbol{R}=\boldsymbol{R}^\star}} \\
&= \begin{bmatrix} \mathcal{D}\big(g(\mathcal{D}(\boldsymbol{R}^\star C(1-l) - \boldsymbol{z})\big) & \mathcal{D}\big(g \circ \boldsymbol{N}^\star\big)(1-l)C \\[1ex] -\mathcal{D}(\boldsymbol{R}^\star)C^T + lD^T\mathcal{D}(\boldsymbol{R}^\star)C^T & -I\tau^{\circ -1}\mathcal{D}(C^T\boldsymbol{N}^\star) + lD^T\mathcal{D}(C^T\boldsymbol{N}^\star) \end{bmatrix}
\end{aligned}
$$

Here, $I$ is the identity matrix. After each assembly, $J$ was evaluated at the equilibrium abundances and its eigenvalues calculated. We find that the real part of the dominant eigenvalue (right-most on the real axis; e.g., Fig S3) of all assembled parent as well as coalesced communities is negative and real. That is, feasibility guarantees local asymptotic stability in our model systems. Note that here, leakage $l$ is not a vector because during assembly, we assumed that all species had the same community-level leakage.

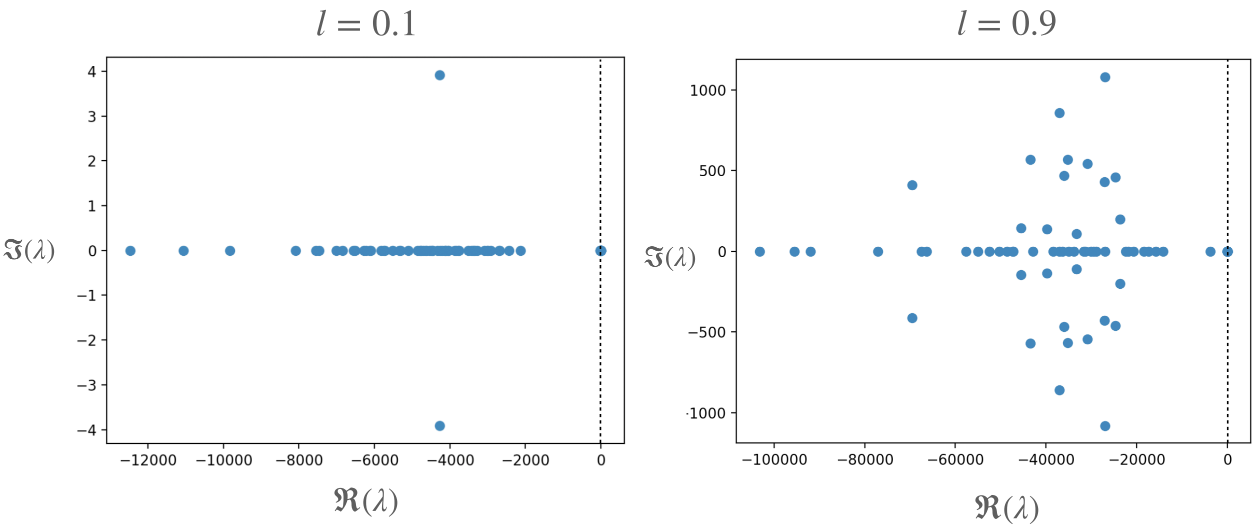

Figure S3: Real (x axis) and complex (y axis) parts of eigenvalues of J when evaluated at equilibrium for two example communities with $l = 0.1$ and $l = 0.9$. In both cases, all eigenvalues are negative (with the leading eigenvalues very close to zero), indicating local stability of the equilibrated communities.

#### S1.4 Relationship with other microbial consumer-resource models

Here we describe how our model is related to two other recent ones that have been used to study microbial community assembly and coalescence.

The mapping between the notation used in Tikhonov (2016) (T), Marsland et al. (2019) (M) and those used here is provided in the following table:

| Notation for... | M | Here | T |
|---|---|---|---|
| Species index | $i$ | $\alpha$ | $\vec{\sigma}$ |
| Species abundance | $N_i$ | $n_\alpha$ | $n_\sigma$ |
| Resource a species can harvest | $\vec{c_i}$ | $\vec{c_\alpha}$ | $\sigma_i$ |
| Resource supply | $\kappa_\alpha$ | $\kappa_j$ | $R_i$ |
| Dilution rate | $\tau_\alpha$ | $\infty$ | $NA$ |
| Minimal resource requirement (maintenance cost) | $m_i$ | $z_\alpha$ | $\chi_{\vec{\sigma}}$ |
| Resource weight | $w_\alpha$ | $\vec{1}$ | $\vec{1}$ |
| Resource $\rightarrow$ biomass conversion factor | $g_i$ | $g_\alpha$ | $(\tau_0 \chi_{\vec{\sigma}})^{-1}$ |
| Leakage factor | $l_\alpha$ | $l$ | 0 |
| Metabolic matrix | $D_{\alpha\beta}$ | $(D_{kj})^T$ | NA |

#### With Marsland et al. (2019)'s Model

Our model differs from the version used in Marsland et al. (2019) in the following respects: (i) all resources contain the same amount of energy (taken to be 1 for simplicity), (ii) a type I functional response, (iii) binary consumer preferences, (iv) a shared core metabolism encoded in $D$, (v) a common leakage fractions for all species and resources, and (vi) a complex environment where all resources are externally supplied in equal amounts. We address the implications of these assumptions in the Discussion section.

1. All resources contain the same amount of energy ($\omega_j = 1$).
2. We only consider a type-1 functional response ($c(R_j) = R_j$).
3. Consumer preferences are binary, instead continuously distributed between 0 and 1.
4. We use a different cost function (further details below) from Marsland et al. (2019) who assume that that maintenance cost $z_\alpha$ is a random fixed quantity for each species.

##### With Tikhonov (2016)'s Model

Tikhonov (2016) did not explicitly model resource dynamics, but we can establish the relationship between that model and the one used here as follows. If we take the system of equations in S1 and assume that (i) there is no leakage ($l = 0$) and (ii) the dilution rate is very low, such that $\tau^{-1} \approx 0$, we can make the assumption that molecular (resource) dynamics are faster than population dynamics, and therefore that resource concentration $R_j$ at any moment quickly equilibrates to reflect the instantaneous demand, i.e., $dR/dt \approx 0$. This allows us to separate the time scales between resource and population dynamics, recovering the model used by Tikhonov (2016):

$$\frac{dN_\alpha}{dt} = g_\alpha N_\alpha \Big( \underbrace{\sum_j c_{\alpha j} \frac{\kappa_j}{\sum_\alpha N_\alpha c_{\alpha j}} - z_\alpha}_{\text{Resource surplus } \Delta_\alpha} \Big)$$

$$R_j = \frac{\kappa_j}{\sum_\alpha N_\alpha c_{\alpha j}}$$

Furthermore, we use a different cost function (previous section) than the one used by Tikhonov.

### S2 Modulating net competition levels

We are interested in generating communities spanning a broad range of competition and facilitation levels, i.e., different competition and facilitation levels ($\mathcal{C} - \mathcal{F}$); Main text Eqs 3–6). To this end we sample the $c_{\alpha j}$ and $D_{jk}$ values for each community with certain constraints (Fig S4).

#### S2.1 Modulating competition level

First, consider the problem of increasing the competition level ($\mathcal{C}$) of the community. The species $\alpha$ has a binary vector $\vec{c_\alpha}$ of length $m$ that specifies if resource $j$ is used ($c_{\alpha j} = 1$) or not ($c_{\alpha j} = 0$). We sample the resource preference vector of each consumer sequentially (that is, one species at a time), in a way that allows us to modify the niche similarity between them. For each consumer $\alpha$, the sampling probability of each resource $j$, $p_{\alpha j}$, is re-evaluated (as in a preferential attachment process, Barabási & Albert (1999)) such that those resources

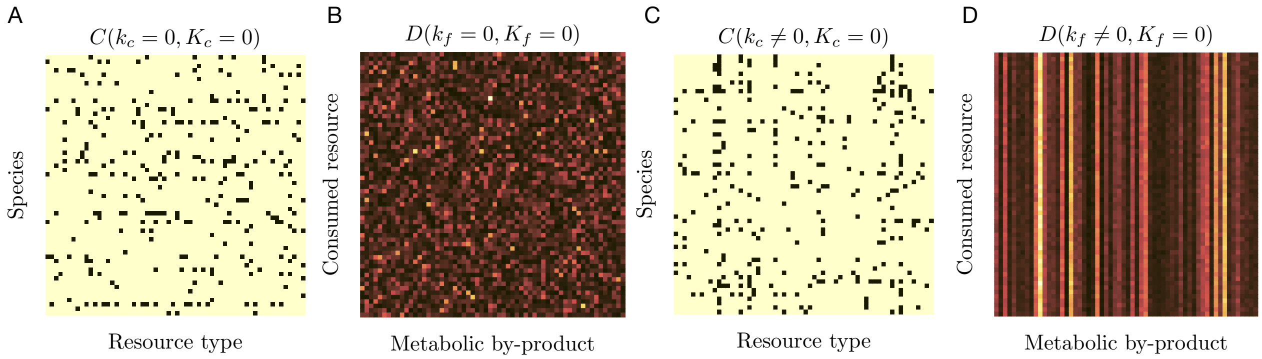

Figure S4: **Examples of differently-structured preference ($C$) and metabolic ($D$) matrices**. These have been generated with different combinations of the competition and facilitation factors $k_c$, $k_f$ in systems of 60 resource types and 60 consumer species Here, $K_c$ and $K_f$ are set to zero. Increasing these values would move the regime towards more structured resource use (See section S2). **A & B**: Uniformly random matrices, where all four parameters are 0. **C & D**: As $k_c$ and $k_f$ are increased the regime moves towards greater preferential feeding, where more demanded resources are more likely to be consumed (increase of $k_c$), but also secreted at higher fractions (increase of $k_f$). In the metabolic matrices (B & D), lighter colours indicate higher values of resource fractions secreted.

that have been frequently sampled by previous species receive a higher probability in the current species.

More specifically, we construct the the resource preference matrix $C$ by sampling these rows $\vec{c_\alpha}$'s as follows. First, we sample $m_\alpha$, the number of resources that species $\alpha$ uses, from an exponential distribution. This choice is supported by experimental evidence (Sung et al. 2017). Second, to determine which resources are used by species $\alpha$, we sample $m_\alpha$ resources with probability vector $\vec{p_\alpha}$. Note that in this sampling scheme, 'iteration number' and 'species' are equivalent, and denoted by index $\alpha$.

For species $\alpha = 1$ all resources have the same probability of being sampled $1/m$, where $m$ is the total number of resources. This assumption is consequent with the absence of a resource hierarchy, since they all carry the same energy. After each iteration, the sampling probability of each resource changes according to what has been sampled previously. Let $d_{\alpha j}$, denote the cumulative demand of resource $j$ when the metabolic preferences of species $\alpha$ are sampled. That is, the number of consumers of resource $j$ at iteration $\alpha$.

$$d_{\alpha j} = \sum_{i=1}^{\alpha} c_{ij}.$$

Based on $d_{\alpha j}$, we then compute the probability that species $\alpha$ is assigned resource $j$ as one of its preferences

$$p_{\alpha j} = (1 - k_c)\frac{1}{m} + k_c \frac{d_{\alpha - 1 j}}{\sum_j d_{\alpha - 1 j}}, \tag{S6}$$

where the denominator represents the total number of preferences sampled up until iteration $\alpha - 1$, and acts as a normalization constant, and together with the numerator represents the normalized cumulative demand ,

$$\tilde{d}_{\alpha-1j} = \frac{d_{\alpha-1j}}{\sum_j d_{\alpha-1j}}.$$

The strength with which $p_{\alpha j}$ deviates from a uniform distribution is given by the parameter $k_c \in [0, 1)$ that is, how much consumers prefer highly-demanded resources, such that when $k_c = 0$ the sampling is uniformly random (Fig S4A); and as $k_c \to 1$ the feeding becomes increasingly preferential (Fig S4C). Pseudocode for the metabolic preferences sampling procedure is given in **Procedure 1**.

**Procedure 1:** Sampling of metabolic preferences

```
for α ∈ {1, . . . , s} do
    Sample m_α from an exponential distribution
    Sample vector v⃗ of m_α integers ∈ {1, . . . , m} with probability vector p⃗(α)
    Switch on sampled preferences c⃗_α[v⃗] = 1
    Update d⃗_α
    Update p⃗_α using the new d⃗_α
end
```

#### S2.2 Modulating facilitation level

Effective competition can alternatively be reduced through the indirect positive effect of facilitation, that is, by increasing $\mathcal{F}/l$. This can be achieved if resources that are highly demanded are also secreted in larger fractions (Fig S4E). For this we need to modulate the structure of the the metabolic matrix $D$. Each element of this matrix, $D_{jk}$, specifies the fraction of leaked energy from resource $j$ that is released in the form of resource $k$. Note that by definition $\sum_j D_{jk} = 1$. Thus, we sample each row of the metabolic matrix $D$ (Fig S4D) from a Dirichlet distribution with a specific concentration parameters $q_{jk}$. The elements of $q_{jk}$ increase proportionally with the demand of each resource $d_j$ and the cooperation factor $k_f$ as

$$q_{jk} = \frac{1}{u}(1 + k_f d_j). \tag{S7}$$

Thus, $k_f$ sets the degree of structure of $D$. When $k_f \to 0$ the metabolic matrix has no structure; all elements of the concentration parameter are the same (flat Dirichlet distribution) and therefore, all resources are released at equi-probable fractions. As $k_f \to 1$ the structure of $D$ becomes fully determined by the resource demands of the community, so that more demanded resources are released at higher fractions. The factor $u$ in Eq S7 controls the sparsity of the metabolic network, ranging from a fully connected network when $u \to 0$ to a sparse one-to-one network when $u \to 1$.

Note that although the above two methods for sampling the elements of $C$ and $D$ share similarities, they are conceptually different. First, the sampling of $C$'s elements is fully random, in the sense that a vector of probabilities is constructed first, and then preferences randomly sampled from it. On the other hand, the sampling of $D$'s elements has a random term.

### S3 Further details of the community coalescence simulations

#### S3.1 Coalescence simulation procedures

##### Recursive coalescence

Given two microbial communities $A$ and $B$, and two community leakage values $l_A$ and $l_B$, such that we implement the following computational procedure:

1. Assemble community $A$ with leakage value $l_{Ai}$.
2. Assemble community $B$ with leakage values $l_{Bj}$.
3. Perform coalescence event between A and B and record parent community dominance (defined below)
4. Change the value $l_{Bi} \rightarrow l_{Bi+1}$, and repeat steps 2 and 3 until vector $l_B$ has been fully traversed.
5. Replicate steps 1-4 in order to gain statistical power.
6. Change value of $l_{Aj} \rightarrow l_{Aj+1}$ and repeat steps 1-5 until vector $l_A$ has been fully traversed.

Pseudocode for the this coalescence simulation is shown in **Procedure 2**. Note that our

---

**Procedure 2:** Recursive coalescence

---

```
begin
    for 1 ≤ i ≤ length(l⃗_A) do
        while better statistical power is required do
            for 1 ≤ j ≤ length(l⃗_B) do
                Assemble community A with leakage value l_{A_i}
                Assemble community B with leakage value l_{B_j}
                Coalesce communities A & B
                Record parent community dominance
```

---

recursive coalescence procedure does not guarantee that every time parent community $B$ is re-assembled (step 2), the present species and their abundances will remain the same as those in the community with the previous leakage value. Thus, at each re-assembly of community B, we check that changing the leakage does not affect significantly the community composition. With this purpose, we calculate the auto-correlation between community abundance vectors of consecutive iterations, and find that it remains close to 1 along all the studied leakage range (see Fig S5).

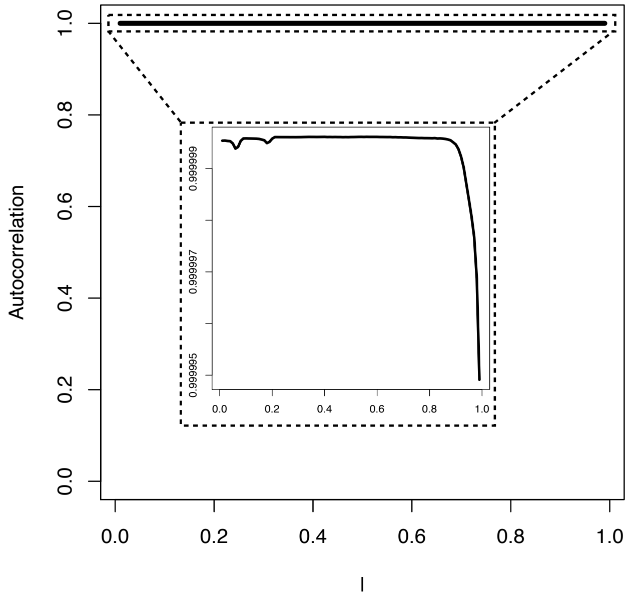

Figure S5: For recursive coalescence, auto-correlation of vector of species abundances in community B for consecutive re-assemblies of this community, along the studied leakage range. (inset) Zoom in to show that, even though the auto-correlation remains consistently $\approx 1$, it starts decreasing from a certain leakage value. While the reason for this decrease is interesting and a potential line of future work, it does not affect our results because of the very small change of magnitude in autocorrelation values over this range.

##### Serial coalescence

Given two sets of *target* and *invader* communities (each of the latter with different leakage values) we implement the following computational steps:

1. Assemble target community $T_0$ with leakage $l_{Ti}$.
2. Assemble a random invader community $W$ with leakage $l_{Wj}$.
3. Coalesce $T_0 + W_1 = T_1$, and record properties of community $T_1$.
4. Repeat steps 2 and 3 until community $T_n = T_{n-1} + W_n$ is reached.
5. Change leakage of the $W$ communities $l_{Wj} \rightarrow l_{Wj+1}$.
6. Repeat steps 2-5 until leakage vector $\vec{l_W}$ has been fully traversed.
7. Change leakage of $T$ communities $l_{Ti} \rightarrow l_{Ti+1}$.
8. Repeat steps 1-7 until vector $\vec{l_T}$ has been fully traversed.
9. Replicate steps 1-8 in order to gain statistical power.

Pseudocode for the serial coalescence simulation is shown in **Procedure 3**.

**Procedure 3:** Serial coalescence

```
Procedure begin
    while better statistical power is required do
        for 1 ≤ i ≤ length(l_T) do
            Assemble target community T_0 with leakage l_{T_i}
            for 1 ≤ j ≤ length(l_W) do
                for 1 ≤ k ≤ n do
                    Assemble invader community W_k with leakage l_{W_j}
                    Coalesce T_{k-1} and W_k to form target community T_k
                    Record properties of target community T_k
```

### S4 Adding consumer guild structure

Recent empirical studies suggests that microbial species tend to form guilds with similar metabolic capabilities, thus introducing some degree of functional redundancy in the communities they form Louca et al. (2018), Enke et al. (2019). Theoretical studies support these observations Goldford et al. (2018), Marsland et al. (2020), Fant et al. (2021). We therefore add further structure to the matrices $C$ and $D$, by partitioning resources into classes, and constraining consumers to feed on a preferred class, but leak to any other, forming consumer guilds. Adding this structure yields two interaction layers (imagine superimposing Figs S6B and 2C with Figs S6A and 2D): inter-guild facilitation and competition between consumers preferring distinct resource classes, and intra-guild facilitation and competition, which stems from the previously-imposed preferential feeding, yielding an effective secretion matrix like the one plotted in Fig S6C.

Resource preferences in this scenario are assigned similarly to the unstructured preferential feeding described in Section S2, except that the probability that species $\alpha_A$ (which feeds preferentially on resource class $A$) samples resource $j$, is now weighted up or down depending on whether $j$ belongs in guild $A$, or not, respectively (Fig S6B). To this end, we define the form of this function to be:

$$p^A_{\alpha j} = \begin{cases} M\left((1-k_c)\dfrac{1}{m} + k_c\dfrac{d_{\alpha-1j}}{\sum_j d_{\alpha-1j}}\right)(1+K_c) & \text{if } j \in A \\ \dfrac{N}{m-n_c}(1-K_c) & \text{otherwise,} \end{cases} \tag{S8}$$

where $M$ and $N$ are normalization constants that ensure $\sum_j p_{\alpha j} = 1$. Note that $K_c$ modulates the amount of structure in the matrix, ranging from no structure when $K_c = 0$ to maximum guild structure when $K_c = 1$. In order to obtain expressions for the normalization constants $M$ and $N$ we impose the following constrains on each piece of Eq S8

$$p^1_\alpha = \sum_{C(j)\in T} p_{\alpha j} = \frac{n_c}{m}(1-K_c) + K_c \tag{S9}$$

and

$$p_\alpha^0 = \sum_{C(j)\notin T} p_{\alpha j} = \left(1 - \frac{n_c}{m}\right)(1 - K_c). \tag{S10}$$

These two constraints guarantee that two necessary conditions are satisfied; (1) the total probability sums to one, since $p_\alpha^1 + p_\alpha^0 = 1$, and (2) probability of sampling resources in (outside) the preferred class approaches 1 (0) when $K_c$ increases (decreases). We then solve for $M$ and $N$ by expanding equations S9 and S10 using the definition in expression S8 for $p_{\alpha j}$

$$M\left((1-k_c)\frac{n_c}{m} + k_c\frac{T_c}{T}\right)(1+K_c) = \frac{n_c}{m}(1-K_c) + K_c$$

$$M = \frac{K_c + \frac{n_c}{m}(1-K_c)}{(K_c+1)\left(\frac{n_c}{m}(1-k_c) + \frac{T_c}{T}k_c\right)}$$

and

$$N = 1 - \frac{n_c}{m},$$

where $T = \sum_j d_{\alpha-1j}$, and we have used the following expressions

$$\sum_{C(j)\in T} 1 = n_c \qquad \sum_{C(j)\notin T} 1 = m - n_c \qquad \sum_{C(j)\in T} d_{\alpha-1j} = T_c.$$

Thus, the closed form of the sampling probability under the general scenario (Eq 7 main text) is

$$p_{\alpha j} = \begin{cases} \left(K_c + \frac{n_c}{m}(1-K_c)\right)\dfrac{\left((1-k_c)\frac{1}{m} + k_c\frac{d_{\alpha-1j}}{T}\right)}{\left((1-k_c)\frac{n_c}{m} + k_c\frac{T_c}{T}\right)} & \text{if } j \in A \\ \frac{1}{m}(1-K_c) & \text{otherwise.} \end{cases} \tag{S11}$$

The metabolic matrix $D$ (Fig S6A) is constructed such that the fraction of leaked by-product $k$ is lower if it belongs to the same class as the consumed resource $j$ (elements within block-diagonals of $D$), and higher otherwise (off-block diagonal elements of $D$). The prominence of this structure in the matrix is given by the inter-guild facilitation factor $K_f$. Therefore, we sample each row of $D$ from a Dirichlet distribution with concentration parameters $q_{jk}$

$$D_{jk} = \text{Dir}\left(q_{j1}, q_{j2}, \ldots, q_{jm}\right)_k,$$

where the concentration parameter depends on the cumulative demand as specified in the previous section (see Eq S7). Additionally, the value of $q_{jk}$ decreases with the inter-guild facilitation factor $K_f$ if uptaken and leaked resources belong to the same (resource) class, and increases with $K_f$ in the opposite case. With these conditions, the expression for the concentration parameter is

$$q_{jk} = \begin{cases} \dfrac{(1+k_f d_j)}{uM_{C(j)}}(1-K_f) & \text{if } A(j) = A(k) \\ \dfrac{(1+k_f d_j)}{u(M-M_{C(j)})}(1+K_f) & \text{otherwise.} \end{cases} \tag{S12}$$

Here, $u$ is the sparsity of the metabolic network, ranging from a fully connected network when $u \to 0$ to a sparse one-to-one network when $u \to 1$. $M_A$ is the number of consumers in class $A$, to which resource $j$ belongs. Note that in expressions S11 and S12 if we make $K_c = K_f = 0$, all the resources belong to the same class, and we recover equations S6 and S7 from the previous version.

#### S4.1 Coalescence between communities with guild structure

When coalescence is simulated between pairs of communities having guild structure, we find that in the low leakage regime, where significant competition is present, the same result (yellow and red lines in Fig S6D) as in the case of random coalescence with only preferential feeding (Fig S2C) is recovered. We find a positive correlation even in the high leakage regime where facilitation is of the same order of magnitude as competition, indicating that our results qualitatively hold for structured communities as well. Note that competition between guilds, and facilitation within guilds, are both very weak. Therefore, here we calculated community level competition as the average over the block diagonal elements, and community level facilitation as the average over the off-block diagonal elements.

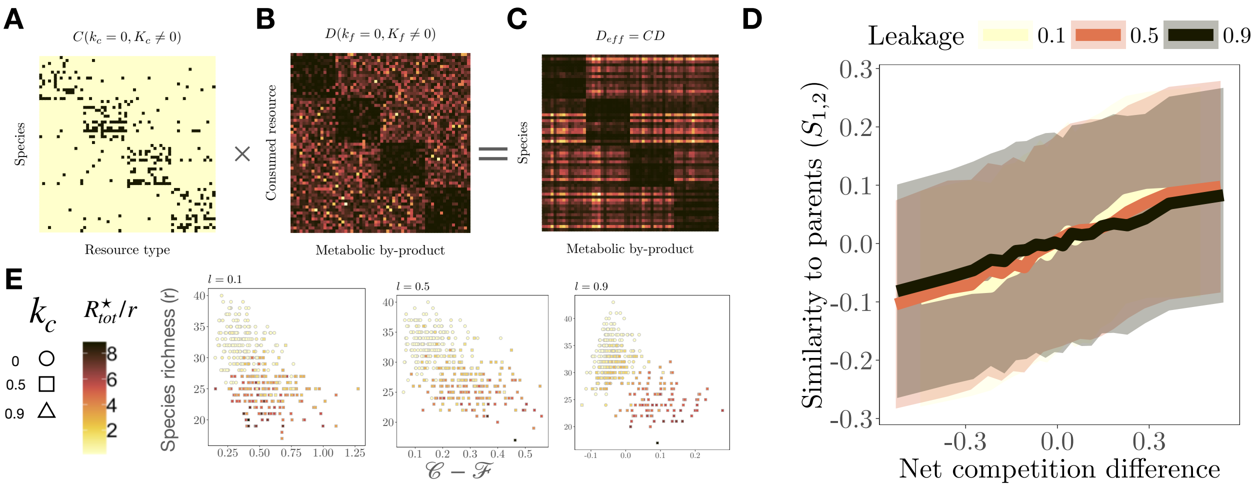

Figure S6: **Community coalescence with consumer guilds present.** **A-C**: Example of a metabolic matrix, preference matrix, and effective secretion matrix, with consumer guild structure. **D**: Similarity to the parent community as function of the binned mean (20 bins) over parent communities with similar difference in competition levels $\mathcal{C}_1 - \mathcal{C}_2$ (solid line) $\pm$ 1 standard deviation (shaded) for the three leakage values. The post-coalescence community is more similar to its less competitive parent. **E**: Species richness ($r$) as a function of community-level net competition ($\mathcal{C} - \mathcal{F}$), coloured by total resource concentration reached at steady state ($R^\star_{tot}$). The observed negative correlation for all values of leakage confirms that less competitive communities are more species-rich, and deplete resources more efficiently (brighter colours, which correspond with lower levels of $R^\star_{tot}$) are scattered towards the top left of the plot.
